# Controlling human stem cell-derived islet composition using magnetic sorting

**DOI:** 10.1101/2024.11.19.624394

**Authors:** Allison B. Kelley, Mira Shunkarova, Marlie M. Maestas, Sarah E. Gale, Nathaniel J. Hogrebe, Jeffrey R. Millman

**Author notes:** To whom correspondence should be addressed: Nathaniel J. Hogrebe,; Jeffrey R. Millman.

## Abstract

Stem cell-derived islets (SC-islets) consists of multiple hormone-producing cell types and offer a promising therapeutic avenue for treating type 1 diabetes (T1D). Currently, the composition of cell types generated within these SC-islets currently cannot be controlled via soluble factors during this differentiation process and consist of off-target cell types. In this study, we devised a magnetic-activated cell sorting (MACS) protocol to enrich SC-islets for CD49a, a marker associated with functional insulin-producing β cells. SC-islets were generated from human pluripotent stem cells (hPSCs) using an adherent differentiation protocol and then sorted and aggregated into islet-like clusters to produce CD49a-enriched, CD49a-depleted, and unsorted SC-islets. Single-cell RNA sequencing (scRNA-seq) and immunostaining revealed that CD49a-enriched SC-islets had higher proportions of β cells and improved transcriptional identity compared to other cell types. Functional assays demonstrated that CD49a-enriched SC-islets exhibited enhanced glucose-stimulated insulin secretion both *in vitro* and *in vivo* following transplantation into diabetic mice. These findings suggest that CD49a-based sorting significantly improves β cell identity and the overall function of SC-islets, improving their effectiveness for T1D cell replacement therapies.

## INTRODUCTION

More than 60,000 people in the United States are diagnosed every year. This currently impacts approximately 1.45 million people and is projected to increase to 2.1 million people by 2040^1^. During T1D, the autoimmune system selectively destroys the endogenous insulin-producing pancreatic β cells within the islets of Langerhans, leading to loss of glucose homeostasis. Current standard of care focuses on accurately administering exogenous insulin and diligently monitoring blood glucose levels. This treatment emphasizes the importance of glycemic regulation but fails to precisely mimic the tight regulation of blood glucose levels achieved by primary islets, leading to long-term complications arising from elevated blood glucose levels over the lifetime of the patient.

To address these inefficiencies in glycemic regulation by exogenous insulin therapy and ease the burden of diabetes management, methods have been developed in recent years to generate SC-islets containing stem cell-derived β cells (SC-β cells) to be used as a cell replacement therapy^2^. While the latest protocol iterations have improved SC-islet function and maturation, these cells still function less efficiently than primary human islets, prompting further investigation into the optimization of the SC-islet cell population. Concurrently, transplantation of cadaveric human islets has been shown to restore normal glycemic regulation, providing strong evidence that replacing lost β cell mass is an effective strategy for resolving T1D. Consequently, much research is now focused on improving the generation, identity, and function of SC-islets as well as methods for effective transplantation for that requires minimal or no immunosuppressant intervention^3, 4^.

While current SC-islet differentiation protocols are quite effective at generating endocrine cell types from hPSCs, they generate multiple cell populations in an imprecise manner^5, 6^, which includes expected pancreatic islet cell populations (β, α, and δ cells) but also undesired off-targets, such as seemingly enterochromaffin cells that normally develop in the intestines^7–9^. Some of these issues have been improved with protocol refinements, SC-islet aggregation techniques, and modulation of additional pathways important in lineage determination^10–15^. Despite this progress, precisely controlling the composition of cell types within these SC-islets to optimize their function has been difficult, which is further exacerbated from the heterogeneity observed across differentiation protocols and cell lines^2, 10^.

Cell sorting is a potential method to improve SC-islet differentiation. CD142, CD200, CD318, GP2, CD177, and custom monoclonal antibodies have been used to sort early endodermal and pancreatic progenitor cells to help with subsequent differentiation in later stages.^16–19^. Insulin-positive cells marked with a GFP reporter combined with cluster reaggregation were used to generate SC-islet clusters enriched in SC-β cells that exhibit improved function^14^. Interestingly, CD49a (Integrin alpha-1, ITGA1) is an integrin involved in adhesion with the extracellular matrix and has been identified to be enriched in SC-β cells within SC-islets^7^. Sorting based on CD49a has been used to increase the fraction of SC-β cells and improve function both *in vitro* and *in vivo*^7, 20^. However, given the dynamics of integrin expression across different cell types and culture methodolgies^21^, the applicability of this approach to other differentiation protocols remains unclear, particularly those that occur primarily in adherent cell culture.

Here, we developed a MACS approach to create CD49a-enriched and de-enriched SC-islet populations generated with our adherent differentiation protocol that leverages direct chemical modulation of the actin cytoskeleton^10, 11^. We found that CD49a MACS produces SC-islets with much better control over the ratios of β, α, δ, and enterochromaffin cells. Furthermore, changing SC-islet cluster composition alters the transcriptional identities of cell types, as assessed with single-cell RNA sequencing (scRNA-seq). Importantly, CD49a-enriched SC-islets demonstrated improved function both *in vitro* and *in vivo*. Overall, our findings demonstrate that controlling SC-islet cell composition with CD49a sorting enhances both the identity and function of the β cells, improving their potential effectiveness for a cell replacement therapy for treating T1D.

## RESULTS

### CD49a MACS Modulates SC-Islet Composition

To perform this study, we generated SC-islets from the HUES8 hPSC line using a 6-stage method that we have previously reported.^10, 11^ Here, we devised a method to modulate the cell composition of these SC-islet clusters based on CD49a sorting (**Fig. 1a**). This protocol consisted of single-cell dispersing clusters and immunostaining this cellular suspension for the surface marker CD49a. Using MACS, we created three sorted cell populations: CD49a enriched cells (CD49a+), CD49a de-enriched cells (CD49a-), and cells run through the MACS columns without separation (unsorted controls). After MACS, the resulting cells were cultured in suspension on an orbital shaker for seven days, for which all three conditions successfully formed into three-dimensional cellular aggregates resembling human islets (**Fig. 1b**). To assess the influence of CD49a MACS on SC-islet cellular composition, we single-cell dispersed CD49+, CD49-, and unsorted control clusters, plated them into multiwell plates, immunostained them for markers of the major expected cell types, and quantified the resulting cell populations (**Fig. 1c-d**). We observed the presence of all the major expected cell populations in all three MACS conditions^7, 22, 23^, including β cells (C-Peptide+), α cells (glucagon+ (GCG)), δ cells (somatostatin+ (SST)), and enterochromaffin cells (SLC18A1+). We confirmed that the CD49a MACS affected the composition of the final SC-islets. CD49a+ cells were enriched in β cells while also being de-enriched in α cells and enterochromaffin cells compared to unsorted controls. The opposite trend was observed with CD49a- cells, with CD49a- cells being de-enriched for β cells and enriched for enterochromaffin cells compared to unsorted controls. Compared to CD49a- cells, CD49a+ cells had 4.0±0.4 times higher fraction of C-peptide+ but only 0.38±0.03, 0.48±0.09, and 0.26±0.05 times the fraction of GCG+, SST+, and SLC18A1+ cells. Taken together, we have provided a detailed method for CD49a MACS of SC-islets that can modulate the final composition of the major endocrine cell populations.

**Figure 1.**
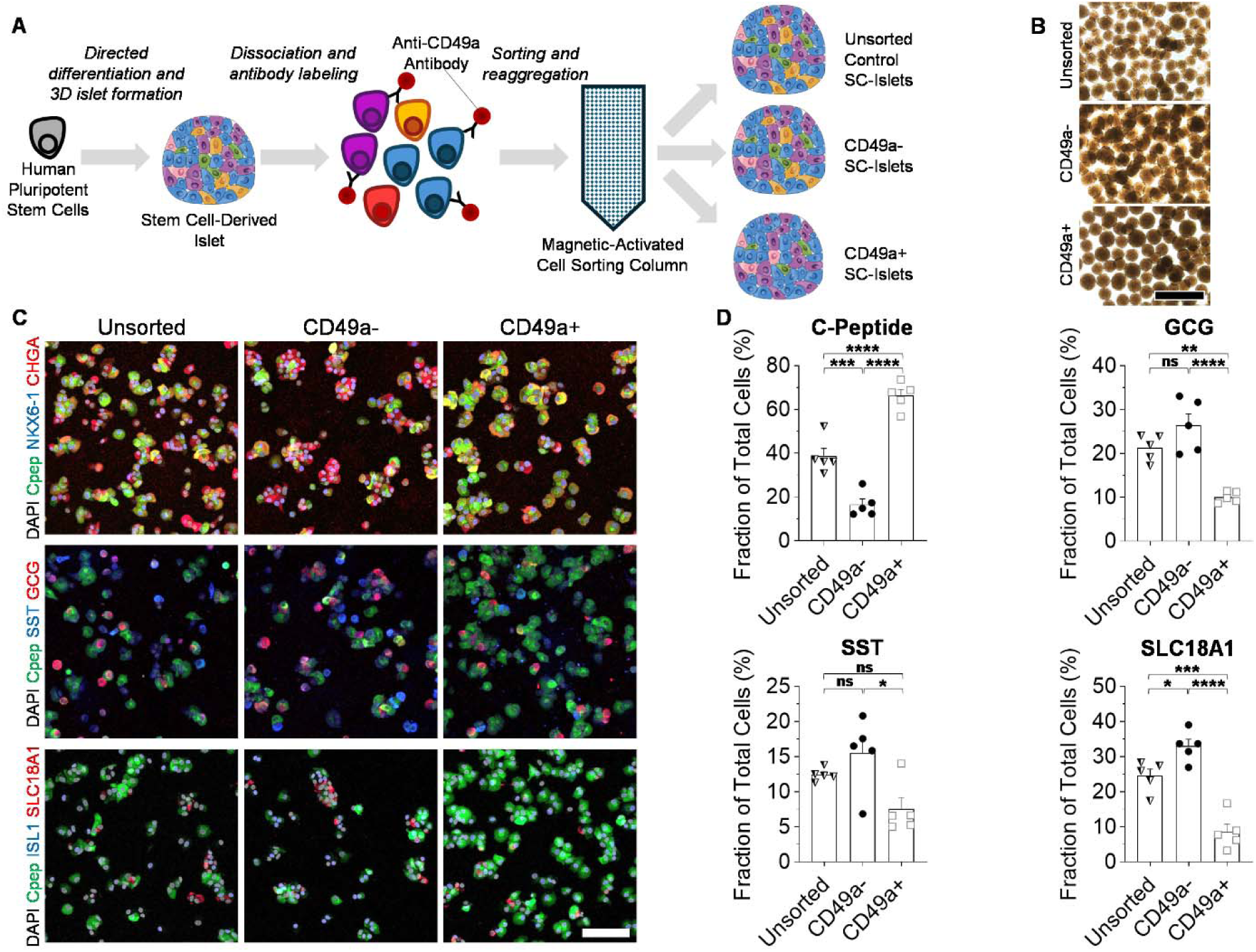
Compositional analysis of SC-islets. (A) Schematic for microscopy analysis of SC-islets after CD49a MACS. (B) Brightfield images of SC-islet clusters 7 days after MACS. Scale bar = 500 µm. (C) Immunofluorescent staining of SC-islets 7 days after MACS single-cell dispersed and plated for 24 hours. Cpep=C-peptide, GCG=Glucagon, SST=Somatostatin. Scale bar = 100 µm. (D) Quantification of the fraction of cells expressing markers of β cells (C-peptide), α cells (GCG), δ cells (SST), and enterochromaffin cells (SLC18A1) (n=5). *p<0.05, **p<0.01, ***p<0.001, ****p<0.0001, and not significant (ns) by two-way unpaired t-test. Bar graphs represent mean, error bars represent s.e.m., and symbols represent individual data points.

### CD49a MACS Improves Cellular Identity of SC-Islets

To better understand how CD49a MACS affects SC-islet identity, we performed scRNA-seq of the resulting cell populations (**Fig. 2a**). We were able to integrate sequencing datasets for CD49a+, CD49a-, and unsorted control SC-islets after MACS and identify major SC-islet populations based on key cellular identity markers (**Fig. 2b-c**)^7, 8, 23–26^. From this integrated dataset, we confirmed that β cells are not only enriched in key identity markers, such as *INS*, *IAPP*, *PDX1*, and *NKX6-1*, compared to other cell types but also that β cells have increased *ITGA1* (CD49a) expression (**Fig. 2d**). We observed all cell populations expressing high levels of *CHGA*, consistent with prior observations that *in vitro* differentiation produces cellular populations that are of high endocrine cell purity.^12^ Furthermore, other major cell types expressed expected cell markers, such as *GCG* and *ARX* for α cells, *SST* and *HHEX* for δ cells, and *TPH1* and *SLC18A1* for enterochromaffin cells^8^. Cross-referencing identified cell types with their CD49a MACS-state revealed that CD49a+ cells were enriched for β cells and de-enriched for other cell types (**Fig. 2e**), consistent with our earlier immunostaining analysis (**Fig. 1c**).

**Figure 2.**
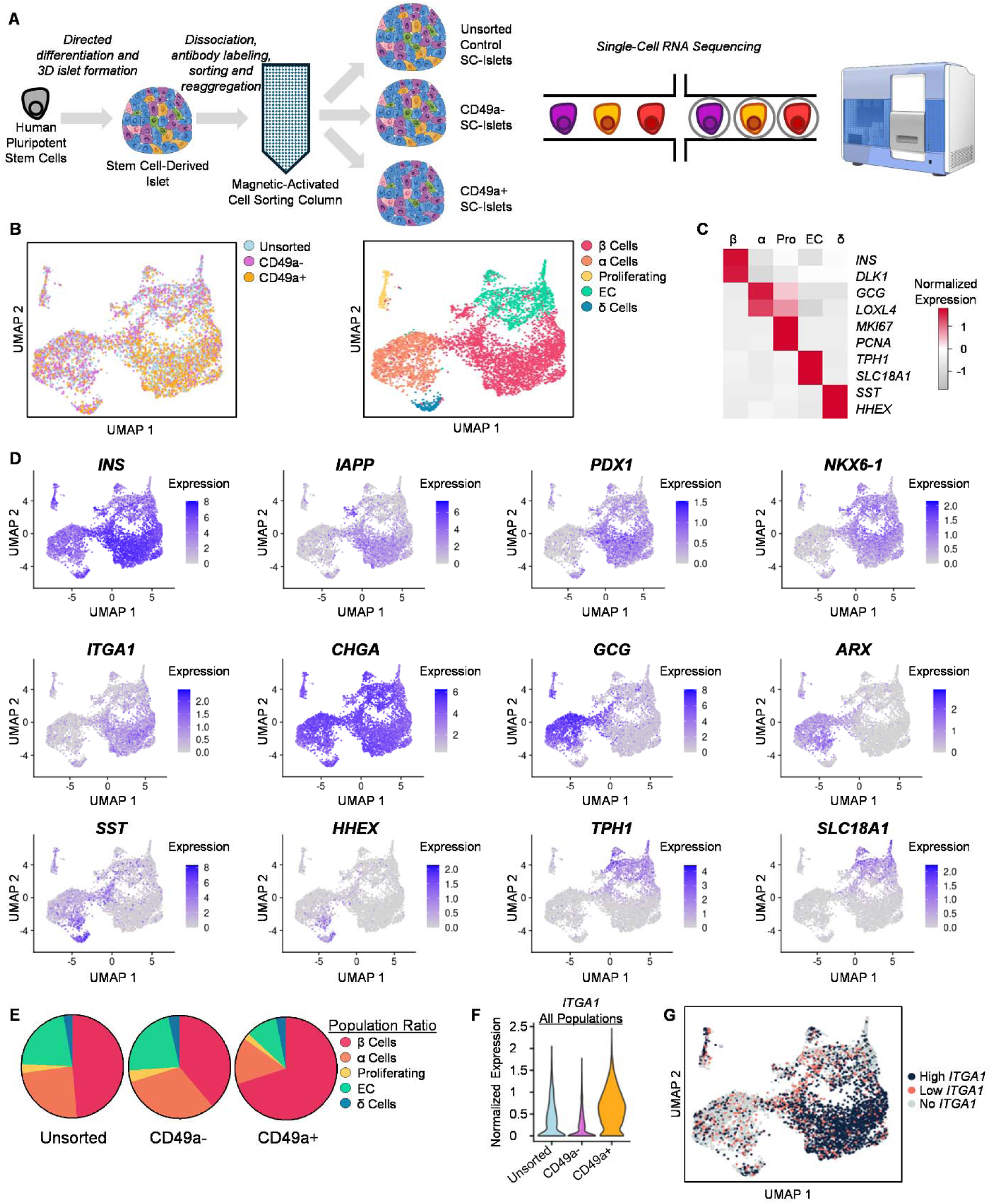
ScRNA-seq of SC-islets. **(A) Schematic for scRNA-seq analysis of SC-islets after CD49a MACS**. (B) UMAP showing MACS condition (left) and islet cell populations (right) based on RNA expression. (C) Heatmap of key markers used to identify islet cell types in the UMAP. (D) UMAP of key markers of interest for SC-islets. (E) Circle graphs showing the fraction of each cell type for each MACS condition. (F) Violin plot showing expression of ITGA1 (CD49a) for all cell populations across each MACS condition. (G) UMAP showing cell populations based on high (top 50%), low (bottom 50%), and no ITGA1 expression.

Furthermore, this bioinformatic analysis revealed that CD49a+ cells were enriched for the relevant transcript *ITGA1* (**Fig. 2f**), confirming the effectiveness of this MACS procedure, and that high *ITGA1* transcript was observed specifically in the β cell population (**Fig. 2g**), as previously reported^7^. To further expand this scRNA-seq analysis, we performed differential expressed gene (DEG) analysis on the subsetted β cell, α cell, δ cell, and enterochromaffin cell populations to (**Fig. 3a; Supplemental Tables 1-4**). We observed that the β cells in the CD49a+ population expressed higher *INS* levels and lower expression of several off-target markers, such as *GCG*, *FEV*, *TPH1*, and *ARX* (**Fig 3b**). We observed this trend in the other key endocrine cell types as well, with the expression of genes associated with β cells such as *INS* and *IAPP* being higher in the CD49a+ population for each cell type. Taken together, this data supports that CD49a MACS results in not only SC-islets of differing composition but also that the identity of these cell types differs after MACS, with CD49a+ being associated with cells with higher β cell identity markers. Most interesting, the β cells from the CD49a+ population appear to partially overcome the shortcomings in immature cellular identity previously identified^3, 4^.

**Figure 3.**
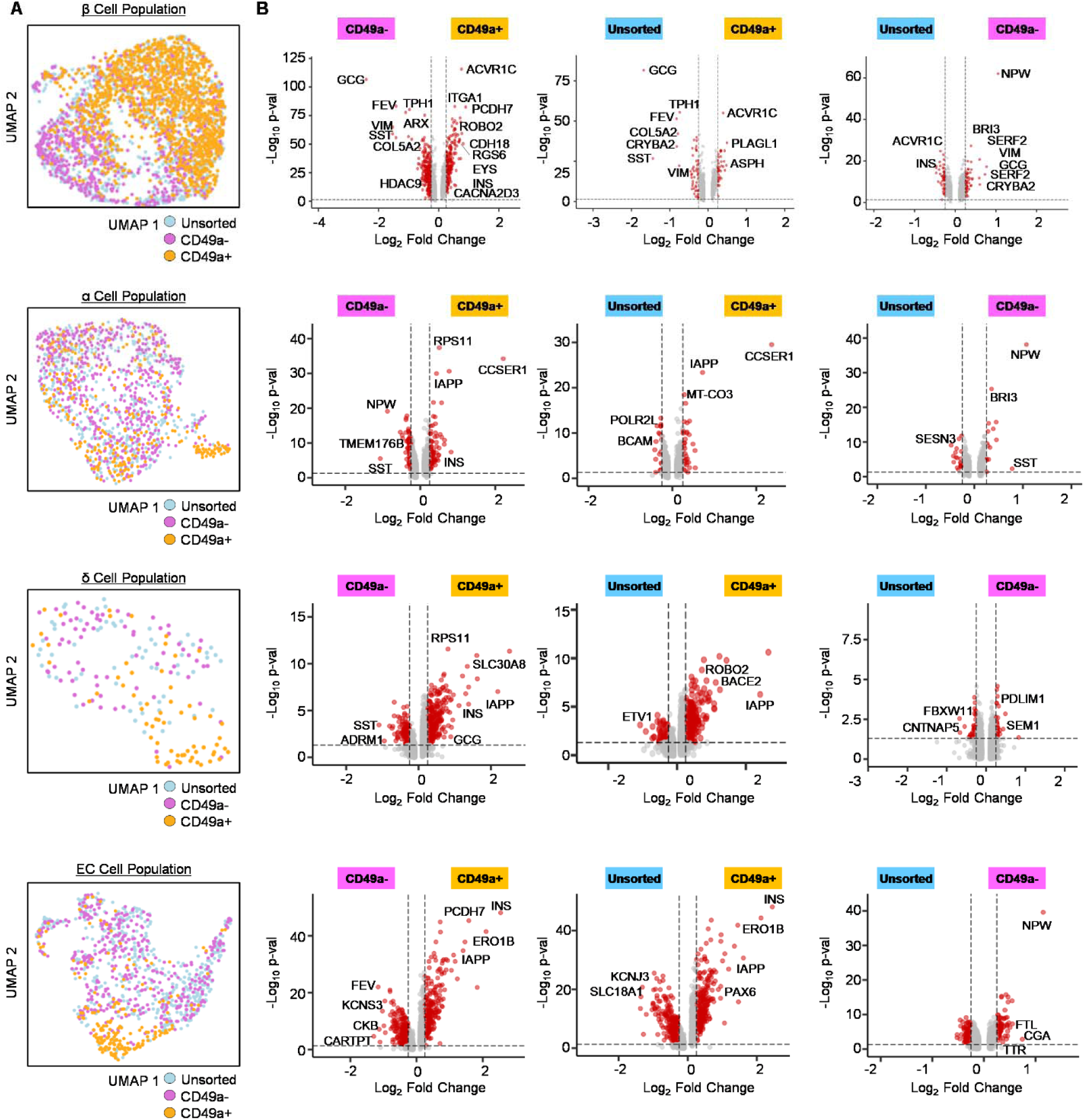
Differential gene expression analysis of SC-islets. (A) UMAP of only β, α, δ, and enterochromaffin cells subsetted from the rest of the cells. (B) Volcano plots showing differential gene expression of subsetted β, α, δ, and enterochromaffin cells for each MACS pair-wise comparison. Thresholds shown are Log_2_ (fold change) >0.25, Log_2_ (fold change) <-0.25, and adjusted *p*-value of <0.05.

### CD49a MACS Improves Function

Our data thus far has indicated large differences in CD49a- cells vs CD49a+ cells. To further examine this in detail, we focused on these two SC-islet populations to evaluate their functional potential (**Fig. 4a**). *In vitro*, we observed that glucose-stimulated insulin secretion was enhanced in CD49a+ SC-islets on both a per cell basis and also on a per β cell basis (**Fig. 4b**). This illustrates that CD49a+ SC-islets have improved function not only because of the increased β cell fraction but also because each β cell is more functional, consistent with our scRNA-seq finding of improved transcriptional identity in β cell of CD49a+ SC-islets (**Fig 3b**). Upon transplantation into non-diabetic NSG mice, we observed significant decreases in blood glucose and improved glucose tolerance for CD49a+ SC-islets vs CD49a- SC-islets but no effect on weight (**Fig. 4c-e**). Furthermore, human C-Peptide levels in the plasma levels were 3.4±0.8 times higher after glucose-stimulation in mice transplanted with CD49a+ compared to CD49a- SC-islets (**Fig. 4f**). While mice with CD49a- SC-islets failed to increase plasma human C-Peptide levels upon glucose injection, mice with CD49a+ SC-islets increased C-Peptide by a factor of 2.5±0.5 after a glucose injection. Immunohistochemical analysis of the grafts confirmed the continued presence of the major SC-islet populations previously identified 10 weeks post-transplantation (**Fig. 4g**). Taken together, this dataset indicates that SC-islets that are enriched for CD49a have enhanced *in vitro* and *in vivo* function, indicating the potential value of such enrichment for diabetes cell replacement therapy.

**Figure 4.**
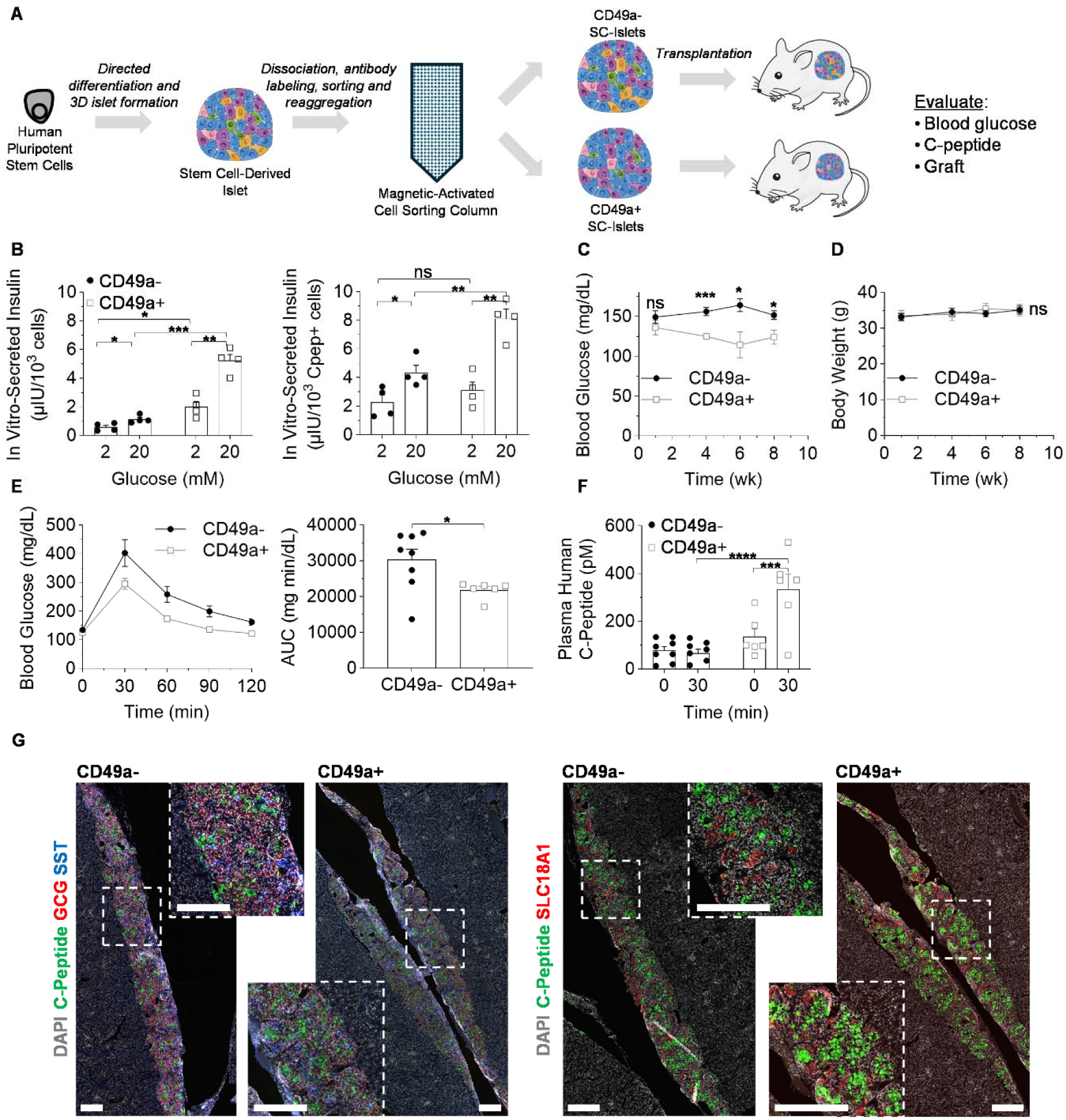
Functional analysis of SC-islets. (A) Schematic for functional analysis of SC-islets after CD49a MACS. (B) *In vitro* glucose-stimulated insulin secretion (n=4). *p<0.05, **p<0.01, ***p<0.001, or not significant (ns) by two-way unpaired t-test. Graph on left normalized by total number of cells and graph on right normalized by number of C-peptide+ cells. (C) NSG mice were transplanted with CD49a- SC-islets (n=6) or CD49a+ SC-islets (n=8) and monitored for up to 8 weeks. Shown are blood glucose measurements. *p<0.05, ***p<0.001, or not significant (ns) by two-way unpaired t-test. (D) Bodyweight of transplanted NSG mice. Not significant (ns) by 2way ANOVA. (E) Glucose tolerance test at 8 weeks after transplantation. Shown are blood glucose concentration over 120 minutes (left) and area under the curve (AUC) calculations. *p<0.05 by two-way unpaired t-test. (F) *In vivo* glucose-stimulated C-peptide secretion before and 30 min after a glucose injection 8 weeks post-transplantation. ***p<0.001 and ****p<0.0001 by two-way unpaired t-test. (G) Immunostaining of SC-islet graft transplanted underneath kidney capsule after 9 weeks. Scale bar=500µm.

## DISUCSSION

In this study, we demonstrated that CD49a-based MACS can successfully enrich SC-islets for insulin-producing β-cells from an adherent-based differentiation protocol while depleting α cells and undesired enterochromaffin cells. Our findings show that CD49a-enriched SC-islets exhibit enhanced β cell identity markers, such as *INS*, reduced identity markers of other endocrine cell types, such as *GCG*, and perform significantly better in glucose-stimulated insulin secretion assays, both *in vitro* and *in vivo*. Importantly, transplantation of CD49a+ SC-islets in mice resulted in improved glycemic control, providing a proof-of-concept for CD49a+ SC-islet enrichment in cell therapy applications.

Previous studies demonstrated that sorting based on CD49a improves SC-islets^7, 20^, and our new study provides additional insights that build upon these data. Prior publications used fully suspension-based differentiation protocols for generating SC-islets^7, 20^. Considering integrin expression is influenced by can be highly influenced by culture environment and cytoskeletal state,^21^ both of which are very different in, and given our mostly-adherent differentiation protocol compared to a fully suspension-based protocol. Furthermore, we previously demonstrated that CD49a/ITGA1 expression changes during SC-islet differentiation.^11^ Thus, we sought to confirm that CD49a sorting would also enhance the effectiveness of our differentiation protocol, which primarily takes place in adherent culture^10, 11^. We found that CD49a enrichment not only increased the proportion of functional β cells but also significantly improved their identity and functionality. Specifically, our scRNA-seq analysis revealed unique transcriptional profiles associated with SC-β cells within the CD49a+ SC-islets. such as the upregulation of on-target markers like *INS* and downregulation of off-target markers such as *GCG* and *FEV*.

Our findings suggest that CD49a enrichment could enhance the therapeutic efficacy of SC-islets for T1D cell replacement therapies^3^. Given that CD49a+ SC-islets demonstrated improved function after transplantation into mice, combining this sorting technique with other technologies, such as encapsulation devices to shield islets from immune attack^27^ or CRISPR-based gene editing to improve immune compatibility^28^, could provide synergistic benefits. Future studies could explore combining CD49a-enriched SC-islets with advanced biomaterials^29^ or other cell types, such as endothelial cells^30^, that promote islet survival and integration *in vivo*.

CD49a-enriched SC-islets also hold promise for *in vitro* disease modeling of diabetes and other metabolic disorders. By providing a more homogeneous population of functional β cells with fewer off-targets, these islets could be used to model disease-specific dysfunctions in patient-derived hPSCs, helping to better understand β cell failure in diabetes. Building on recent advances in microenvironmental and organoid-based *in vitro* disease models^28^, incorporating CD49a-enriched islets into such systems could facilitate the study of β cell-microenvironment interactions under pathological conditions, providing new avenues for therapeutic discovery.

## METHODS

### Stem cell culture

The human embryonic stem cell (hESC) line HUES8 [RRID: CVCL_B207] was used in this study. This was generously provided by Dr. Douglas Melton (Harvard University) under a Material Transfer Agreement (MTA), and we have used this cell line previously^12^. The hESC line were used under the approval of the Washington University Embryonic Stem Cell Research Oversight Committee (approval no. 15-002) with the appropriate consent and conditions. We propagated the undifferentiated stem cells on Matrigel (356230, Corning) coated tissue culture-treated flasks at 100,000 cells/cm^2^ in mTeSR1 (85850, Stemcell Technologies) supplemented with 10 µM Y-27632 (ab120129, Abcam) and cultured in a humidified incubator at 5% CO_2_ and 37°C. The media was replaced every day with new mTeSR1, increasing the volume added each day as the cells grew. After 4 days, the cells were passaged using 0.2 mL TrypLE Express/cm^2^ (12604-039, Gibco) and seeded in mTeSR1 supplemented with 10 µM Y-27632 for either further propagation or differentiation.

### SC-islet differentiation

We differentiated SC-islets using a protocol we previously published^10, 11^. Undifferentiated HUES8 stem cells were seeded onto Matrigel-coated tissue culture-treated 6- well plates or flasks (T75 or T182.5) in mTeSR1 supplemented with 10 µM Y-27632 at a density of 0.63 cells/cm^2^. After 24 hours, differentiation was initiated by replacing the mTeSR1 with Stage 1 media and factors. The cells were fed every day with the following base media and differentiation factors.

#### Base media formulations

<u>Stage 1</u>: 500 ml MCDB 131 (10372019, Gibco) supplemented with 0.8 g glucose (G7528, MilliporeSigma), 0.587 g sodium bicarbonate (S5761, MilliporeSigma), 0.5 g BSA (68700, Proliant Biologicals) and 5 mL GlutaMAX (35050-079, Gibco). <u>Stage 2</u>: 500 mL MCDB 131 supplemented with 0.4 g glucose, 0.587 g sodium bicarbonate, 0.5 g BSA, 5 mL GlutaMAX and 22 mg vitamin C (A4544, MilliporeSigma). <u>Stages 3 and 4</u>: 500 mL MCDB 131 supplemented with 0.22 g glucose, 0.877 g sodium bicarbonate, 10 g BSA, 2.5 mL ITS-X (51500-056, Gibco), 5 ml GlutaMAX and 22 mg vitamin C. <u>Stage 5</u>: 500 mL MCDB 131 supplemented with 1.8 g glucose, 0.877 g sodium bicarbonate, 10 g BSA, 2.5 ml ITS-X, 5 mL GlutaMAX, 22 mg vitamin C, 5 mL penicillin/streptomycin solution (30-002-CI, Corning) and 5 mg heparin (H3149, MilliporeSigma). <u>Stage 6</u>: 500 mL MCDB 131 supplemented with 0.23 g glucose, 10.5 g BSA, 5.2 ml GlutaMAX, 5.2 mL penicillin/streptomycin solution, 5 mg heparin, 5.2 mL MEM nonessential amino acids (20-025-CI, Corning), 84 μg ZnSO4 (10883, MilliporeSigma), 523 μL Trace Elements A (25-021-CI, Corning) and 523 μL Trace Elements B (25-022-CI, Corning).

#### Differentiation factors

<u>Stage 1 (4 days)</u>: Base medium supplemented with 100 ng/mL Activin A (338-AC, R&D Systems) and 3 μM CHIR99021 (2520691, Peprotech) for the first 24 hours, followed by 3 days of BE1 containing 100 ng/mL Activin A only. <u>Stage 2 (2 days)</u>: Base medium supplemented with 50 ng/mL KGF (AF-100-19, Peprotech). <u>Stage 3 (2 days)</u>: Base medium supplemented with 50 ng/ml KGF, 200 nM LDN193189 (SML0559, MilliporeSigma), 500 nM TPPB (53431, Tocris Bioscience), 2 μM retinoic acid (R2625, MilliporeSigma) and 0.25 μM SANT1 (S4572, MilliporeSigma). <u>Stage 4 (4 days)</u>: Base medium supplemented with 50 ng/mL KGF, 200 nM LDN193189, 500 nM TPPB, 0.1 μM retinoic acid and 0.25 μM SANT1. <u>Stage 5</u> <u>(7 days)</u>: Base medium supplemented with 10 μM ALK5i II (ALX-270-445-M005, Enzo Life Sciences), 1 μM T3 (64245, MilliporeSigma), 1 μM XXI (565790, MilliporeSigma), 0.1 μM retinoic acid, and 0.25 μM SANT1. 1 µM Latrunculin A (10010630, Cayman Chemical) was added to this medium for the first 24 hours only. <u>Stage 6 (>7 days)</u>: Cultures were kept on the plate with the base media for up to 7 days. To generate islet-like clusters, cells were single-cell dispersed with TrypLE and resuspended in 6 mL of ESFM within a 6-well plate at a concentration of 5-8 million cells per well on an orbital shaker (Orbi-Shaker CO_2_, Benchmark Scientific) at 115 RPM.

### Magnetic sorting

Three days after aggregation of SC-islets in stage 6, cell clusters were washed with PBS (Corning; 21-040-CV)) and then single-cell dispersed with prewarmed 37^0^C TrypLE (Life Technologies; 12-604-039) for 10 minutes. The TrypLE was neutralized with phosphate-buffered saline (PBS) (Corning; 21-040-CV), the cells were centrifuged, supernatant was removed, the singularized cells were resuspended in 2% PBS+FBS buffer solution (2% fetal bovine serum (Sigma-Aldrich; F2442-500ML) in phosphate-buffered saline (PBS) (Corning; 21-040-CV)), and counted with Vi-Cell XR (Beckman Coulter). Cells were stained with primary anti-CD49a antibody (BD biosciences; 559594) diluted 1:500 in a 2% PBS+FBS buffer solution for 20 minutes on ice. Cells were washed with a 2% PBS-FBS buffer solution and stained with anti-mouse IgG1 magnetic microbeads (Miltenyi Biotech; 130-047-102) diluted 1:5 in a 2% PBS+FBS buffer solution for 15 minutes on ice. After staining with the magnetic microbeads, cells were washed with a 2% PBS-FBS buffer solution and resuspended in a 0.5% PBS-BSA (2% MACS BSA stock solution (Miltenyi Biotec; 140-001-130.03) in PBS (Corning; 21-040-CV)) buffer solution.

For cell separation, we used a QuadroMACS^TM^ cell separator (Miltenyi Biotec; 130-091-051) with LS Columns (Miltenyi Biotec; 130-042-401) using the positive selection cell separation method. After attaching LS column to the separator, 0.5% PBS-BSA buffer solution was added to the LS column. The cells resuspended in 0.5% PBS-BSA buffer solution were run through the LS column attached to the separator to allow for a CD49a negative fraction to flow through the column. The CD49a positive fraction of cells was washed out from the LS column detached from the separator with the 0.5% PBS-BSA solution by applying pressure with an LS column plunger. Both CD49a negative and CD49a positive fractions were collected and counted. After the number of cells were determined, cells were seeded onto 6-well plates – with ∼5 x 10^6^ in each well – in a stage 6 cell base media cell aggregation for 7 days. Bright field images were taken using a Leica DMi1 inverted light microscope.

### Immunofluorescence staining, imaging, and quantification of cells *in vitro*

Seven days after sorting, SC-islets were single-cell dispersed and plated for 24 hours. To stain cells, we fixed them with 4% paraformaldehyde (PFA) (157-4-100, Electron Microscopy Sciences) at room temperature for 30 minutes. The cells were then blocked and permeabilized for 45 minutes at room temperature with a staining solution consisting of 0.1% Triton X (327371000, Acros Organics) and 5% donkey serum (017000-121, Jackson ImmunoResearch) in PBS (21-040-CV, Corning). These cells were then incubated with primary antibodies diluted in ICC solution overnight at 4 °C, after which the cells were washed with staining solution, incubated with secondary antibodies diluted in staining solution for 2 hours at room temperature, and finally stained with DAPI nuclear stain (D1306, Invitrogen) for 15 minutes at room temperature. Primary antibodies used were: rat anti-C-peptide (GN-ID4-S, DSHB, 1:300), mouse anti-NKX6.1 (F55A12-S, DSHB, 1:100), rabbit anti-CHGA (AB15160, Abcam, 1:1000), mouse anti-somatostatin (NBP2-37447, Novus, 1:500), rabbit anti-glucagon (259A-14, Cell Marque, 1:300), mouse anti-ISL1 (40.2d6-S, DSHB, 1:300), rabbit anti-SLC18A1 (HPA063797, Sigma, 1:500) . The secondary antibodies were used at a 1:300 dilution in ICC. All secondary antibodies were Alexa Fluor raised in donkey: anti-rat 488 (A21208, Invitrogen), anti-mouse 594 (A21203, Invitrogen), anti-rabbit 647 (A31573, Invitrogen), anti-mouse 647 (A31571, Invitrogen), anti-goat 594 (A11058, Invitrogen), anti-rabbit 594 (A21207, Invitrogen). Zeiss Cell Discoverer 7 at the Washington University Center for Cellular Imaging was used for imaging. Quantification was performed by manually counting using Fiji^31^.

### Cell preparation for Single-cell RNA sequencing

SC-islets were single-cell dispersed using TrypLE (ThermoFisher,12604013). The cells were spun down and washed with PBS. Cell counts were obtained for each sample and resuspended at 1,000 cells/uL in DMEM (Fisher, 11-960-044). Live cells were submitted to the Washington University Genome Technology Access Center at the McDonnell Genome Institute. The samples were sequencing using the Illumina NovaSeq X Plus.

### Analysis of Single-cell RNA sequencing

Seurat 4.3.0 was used for the analysis. Quality control on the cells was performed and unsorted cells were filtered for gene numbers between 2,500 and 12,500; CD49a- was filtered between 2,500 and 12,500; and CD49a+ was filtered between 5,000 and 20,000. Cells with high mitochondrial gene expression were excluded from analysis. Each condition was normalized using NormalizeData and the top 2000 variable features were found using FindVariableFeatures. To integrate the three conditions, integration anchors were determined using FindIntegrationAnchors. Next, the anchors found from the previous function were used for the function IntegrateData which will integrate the conditions. After integration, ScaleData, RunPCA, and ElbowPlot were used to determine the dimensions, 1:25, for RunUMAP and FindNeighbors. FindClusters was set to a resolution of 0.4 to determine clustering and dimensional reduction. FindMarkers was used to determine Log_2_FoldChange and P-value which uses the non-parametric Wilcoxon rank sum test.

### *In vitro* glucose-stimulated insulin secretion

To assess the *in vitro* function of these cells, we used culture inserts (PIXP01250, MilliporeSigma) that were placed in a 24-well plate and wetted with 500 µL of Krebs buffer (KrB) (128 mM NaCl, 5 mM KCl, 2.7 mM CaCl_2_, 1.2 mM MgSO_4_, 1 mM Na_2_HPO_4_, 1.2 mM KH_2_PO_4_, 5 mM NaHCO_3_ 10 mM HEPES (15630-080, Gibco) and 0.1% BSA). We collected approximately 50 SC-islet clusters into each insert, washed twice with 1 mL KrB containing 2 mM glucose, transferred the inserts into new wells with 1 mL KrB containing 2 mM glucose, and incubated at 37° C and 5% CO_2_ for 1 hour to equilibrate the samples at low glucose. To transfer each insert between wells, we angled them at approximately 45° until the liquid drained through the membrane from the insert into the well. After 1 hour, we discarded the supernatants and transferred the inserts into new wells with 1 mL KrB containing 2 mM glucose. After incubating for 1 hour, we transferred the inserts into new wells with 1 mL KrB containing 20 mM glucose, saving the supernatants. After an additional hour, we transferred the inserts to new wells containing 1 mL of TrypLE to single-cell disperse the clusters and perform cell counts (Vi-Cell XR, Beckman Coulter). We again saved these supernatants. Taking this together, we quantified the supernatants at low and high glucose with a human insulin ELISA kit (80-INSHU-E10.1, ALPCO), and results were normalized to the cell count for each sample.

### Transplantation studies

All animal studies were performed in accordance with Washington University International Care and Use Committee (IACUC) regulations (protocol #21-0240). We used 7-week-old male immunodeficient mice (NOD.Cg-Prkdc^scid^ Il2rg^tm1Wjl^/SzJ, Jackson Laboratories). Surgery and assessments were performed by unblinded individuals, and mice were randomly assigned to experimental groups. For transplantation, mice were anesthetized with isoflurane and injected with 2×10^6^ SC-islet cells seven days after sorting underneath the kidney capsule. Mice were monitored for up to 8 weeks, periodically monitoring blood glucose and body weight. After 9 weeks, a glucose tolerance test was performed by fasting mice for 4 hours, measuring blood glucose, then injecting with 3 g/kg glucose in 0.9% sterile saline, and blood glucose measurements were made every 30 minutes for up to 2 hours. Furthermore, *in vivo* human glucose-stimulated insulin secretion was measured after 8 weeks by first fasting mice for 4 hours, collecting plasma, then injecting with g/kg glucose injection (G7528, MilliporeSigma) in 0.9% sterile saline, and collecting plasma again after 30 minutes. Human C-peptide from the collected serum was quantified using the Human Ultrasensitive C-Peptide ELISA kit (10-1141- 01, Mercodia). After 10 weeks, mice were sacrificed, and transplanted kidneys collected for immunohistochemistry.

### *In vivo* immunohistochemistry

For histological sectioning, excised mouse kidneys containing transplanted cells were fixed overnight with 4% PFA at 4°C. These samples were submerged in 70% ethanol and sent off to be paraffin embedded and sectioned by Histowiz, Inc (Long Island City, New York). Paraffin was removed from sectioned samples with Histo-Clear (C78-2-G, Thermo Fisher Scientific), and antigen retrieval was carried out in a pressure cooker (2100 Retriever, Electron Microscopy Sciences) with 0.1 M EDTA (AM9261, Ambion). Slides were blocked and permeabilized with ICC solution for 45 min and then incubated with primary antibodies in ICC solution overnight at 4 °C. After 24 hours, the slides were incubated with secondary antibodies for 2 hours at room temperature and then finally sealed with DAPI Fluoromount-G (0100-20, SouthernBiotech).

Primary antibodies used were rat anti-C-peptide (GN-104-S, DSHB, 1:300), mouse anti-somatostatin (NBP2-37447, Novus, 1:500), rabbit anti-glucagon (259A-18, Cell Marque, 1:2), rabbit anti-Caspase 3 (ab32351, Abcam), goat anti-PDX1 (AF2419, R&D, 1:300), mouse anti-CD31 (M0823, Dako, 1:300), rabbit anti-SLC18A1 (HPA063797, Sigma, 1:1000), mouse anti-NKX6-1 (F55A12, DSHB, 1:100), and rabbit anti-Ki67 (ab1667, Abcam, 1:300). The secondary antibodies were Alexa Fluor in donkey and used at a 1:300 dilution in ICC. They were anti-rat 594 (A21209, Invitrogen), anti-mouse 488 (A21202, Invitrogen), anti-rabbit 647 (A31573, Invitrogen), and anti-goat 488 (A11055, Invitrogen). The Zeiss Axioscan 7 Brightfield/Fluorescence Slide Scanner with Colibri 7 LED illumination sources was used for imaging at the Washington University Center for Cellular Imaging.

### Data Availability

The single-cell RNA sequencing data from this study will available at the Gene Expression Omnibus (GEO) under accession number GSE278800. All other data supporting the findings of this study are available from the corresponding author upon reasonable request.

### Code Availability

No custom codes or algorithms were developed for the analysis of this paper. The integration of the datasets and all other analysis was done using Seurat workflow.

### Statistics and reproducibility

Statistical analysis was performed in GraphPad Prism, version 10. The statistical tests used in each panel are listed in the figure captions. P values are indicated as follows: NS, not significant; *p<0.05, **p<0.01, ***p<0.001, ****p<0.0001. All data are presented as the mean, and all error bars represent the standard error of the mean (SEM). The sample size (n) indicated in the figure legends.

## AUTHOR CONTRIBUTIONS

A.K., N.J.H., and J.R.M. designed all experiments. A.K., M.M.M., M.S., S.E.G., and J.R.M. performed all *in vitro* experiments. A.K. and M.S. performed all *in vivo* experiments. M.M.M. performed all computational analysis. All authors wrote, revised, and approved the manuscript.

## ACKNOWLEDGEMENTS

This work was funded by Sana Biotechnology to N.J.H. and J.R.M. Imaging experiments were performed in part through the use of Washington University Center for Cellular Imaging (WUCCI) supported by Washington University School of Medicine, The Children’s Discovery Institute of Washington University and St. Louis Children’s Hospital (CDI-CORE-2015-505 and CDI-CORE-2019-813) and the Foundation for Barnes-Jewish Hospital (3770 and 4642). We thank the Genome Technology Access Center at the McDonnell Genome Institute at Washington University School of Medicine (P30CA91842) and the Washington University Diabetes Research Center (P30DK020579) for additional analysis support. This publication is solely the responsibility of the authors and does not necessarily represent the official view of NIH nor any other funder. Genomesequencer-4 icon by DBCLS (https://togotv.dbcls.jp/en/pics.html) is licensed under CC-BY 4.0 Unported (https://creativecommons.org/licenses/by/4.0/). We thank Erika Brown (Washington University) for careful reading and feedback of this manuscript.

## COMPETING INTERESTS

N.J.H. and J.R.M. are inventors on patents and patent applications related to SC-islets, including intellectual property licensed to Sana Biotechnology. J.R.M. was employed at and has stock in Sana Biotechnology. The remaining authors declare no competing interests.

## Notes

### Competing Interest Statement

This work was funded by Sana Biotechnology to N.J.H. and J.R.M. N.J.H. and J.R.M. are inventors on patents and patent applications related to SC-islets, including intellectual property licensed to Sana Biotechnology. J.R.M. was employed at and has stock in Sana Biotechnology. The remaining authors declare no competing interests.

